# A culture-based approach to study ecological interactions among the microbial species of the human scalp

**DOI:** 10.1101/2023.09.07.556667

**Authors:** Swagatika Bhattacharya, Talia Roth, Suzannah Costa, Ava Santoro, William Mazza, Katharine Z. Coyte, Cécile Clavaud, Kevin R. Foster, Wook Kim

## Abstract

The human scalp hosts an unusually low diversity microbiota dominated by three species: *Cutibacterium acnes*, *Staphylococcus epidermidis*, and *Malassezia restricta*, where characteristic shifts in species’ frequencies are associated with seborrheic dermatitis and dandruff. In order to better understand this important community, here we study the ecological interactions between these scalp species. We establish a new experimental model system that supports the growth of all three species *in vitro* and allows one to selectively enumerate each species from co-culture. Our work reveals the potential for strong ecological interactions within the scalp community. In particular, *C. acnes* greatly benefits from the presence of *M. restricta*, but harms it in return (exploitation), while *S. epidermidis* suppresses both *M. restricta* and *C. acnes*. Our data suggest that the shifts in composition seen in compromised scalps are influenced by ecological interactions between species. We argue that the scalp microbiome should be viewed as an ecological system where species interactions have the potential to contribute to health outcomes.

**Importance:** Our bodies are home to diverse communities of microorganisms, our microbiome, which can be critical for health and wellbeing. The human scalp hosts a relatively simple community dominated by three species: two bacteria, *Cutibacterium acnes* and *Staphylococcus epidermidis*, and one fungus, *Malassezia restricta*. Both dandruff and seborrheic dermatitis are strongly associated with characteristic shifts in the frequencies of these three species. However, how these species affect one another and behave as a community remains poorly understood. Here, we develop a simple experimental system to empirically study how these three species interact and affect one another for the first time. We find that *S. epidermidis* greatly suppresses the growth of the other species, while *C. acnes* specifically exploits *M. restricta*. Our work suggests that the human scalp is an ecological system in which species interactions have the potential to affect health outcomes.

## Introduction

The human body is densely colonized by microorganisms that are present at numbers comparable to human cells (1), and the relative composition of microbial communities vary considerably within and among individuals (2–5). The skin is the largest organ of the human body, and on top of the skin lies a dynamic community of commensal microbes that limit the growth of pathogens (6), such that its disruption has the potential to lead to significant clinical complications (7).

Amongst body sites, the scalp carries a relatively low diversity microbial community (8–10) that is overwhelmingly dominated by three species: the fungus *Malassezia restricta*, and the bacteria *Cutibacterium acnes* and *Staphylococcus epidermidis*. Amplicon and metagenomics studies reveal that *M. restricta* represents 83-97% of all fungal DNA found on the scalp and *C. acnes* and *S. epidermidis* together represent 97-99% of all bacterial DNA, across individuals and geographic regions (11–17). Notably, these studies non-invasively collected samples from the surface of the scalp with a wet swab, suggesting that the three dominant species co-exist on the skin surface. Moreover, a characteristic shift in their frequency is strongly correlated with seborrheic dermatitis and dandruff: increased *M. restricta* and *S. epidermidis*, and reduced *C. acnes* (12, 13). Seborrheic dermatitis and dandruff have long been associated with fungi as antifungal shampoos provide transient relief (18–20). However, the impacts of these shampoos on the wider microbial community of the scalp is poorly described (21) and there remains no single microbial species that is an indicator of dandruff, which hints at a complex causation beyond *Malassezia*. Moreover, *C. acnes*, *S. epidermidis*, and *M. restricta* all have the potential to cause skin infections in particular cutaneous context (22–24), further suggesting that they all need to be considered in the study of scalp biology. Understanding the ecology of the healthy scalp microbiome could help in developing new approaches to maintain scalp health.

We, therefore, decided to study the three dominant scalp species in concert by growing them both alone and together to better understand their ecology. However, growing and studying these diverse natural isolates in the lab poses significant technical challenges as each were thought to have distinct culturing requirements. Here, we present a simple experimental platform to explore the interactions among scalp microbes, which reveals previously unknown relationships between these species.

## Results and Discussion

### Identification of optimal growth parameters for mixed-species culture

A key requirement for a mixed-species experimental system is that each species must grow under a common condition, but there are no known laboratory conditions that commonly support the growth of all three dominant scalp species. The major technical obstacles to overcome were that the three species have disparate oxygen and nutrient requirements: *M. restricta* requires oxygen and lipids, *C. acnes* is an anaerobe that grows on simple carbohydrates and lipids, and *S. epidermidis* is a facultative anaerobe that utilizes diverse carbon sources. Human skin is considered to be hypoxic (1.5-5.0% O_2_) due to the rapid consumption by the epidermis that lack support from the blood vasculature (25). We reasoned that a hypoxic environment may permit the growth of both *M. restricta* and *C. acnes*, and tested strains that were isolated directly from the human scalp in addition to reference type strains. We initially grew all strains on standard media that are routinely used to culture each species (modified Dixon (mDixon) for *M. restricta*, Medium 20 for *C. acnes*, and Tryptic Soy Broth Agar (TSA) for *S. epidermidis*). We chose to focus on solid media instead of liquid to promote structured interactions among the species that also likely take place on the scalp. Inoculated plates were incubated at 32°C to accommodate the growth of *M. restricta* under varying levels of oxygen, and CO_2_ was kept constant at 5% within the hypoxic chamber.

In this survey, *M. restricta* and *C. acnes* strains did not share any common growth conditions while *S. epidermidis* strains grew across all conditions (Table 1). However, we discovered that O_2_ at 2% and CO_2_ at 5% supports the growth of all three species, albeit in different media. We also observed that *M. restricta* strains visibly suffer and fail to produce isolated colonies on mDixon under atmospheric levels of O_2_, unless additional CO_2_ was provided (Fig. 1). A recent study demonstrated that *M. restricta* possesses a carbonic anhydrase, which converts CO_2_ and water to bicarbonate and protons (26). CO_2_ is expected to be produced by respiring scalp cells, and carbonic anhydrases play important roles in pH homeostasis and metabolism across many organisms (27, 28), and we find that it is particularly important for *M. restricta*.

**Figure 1.**
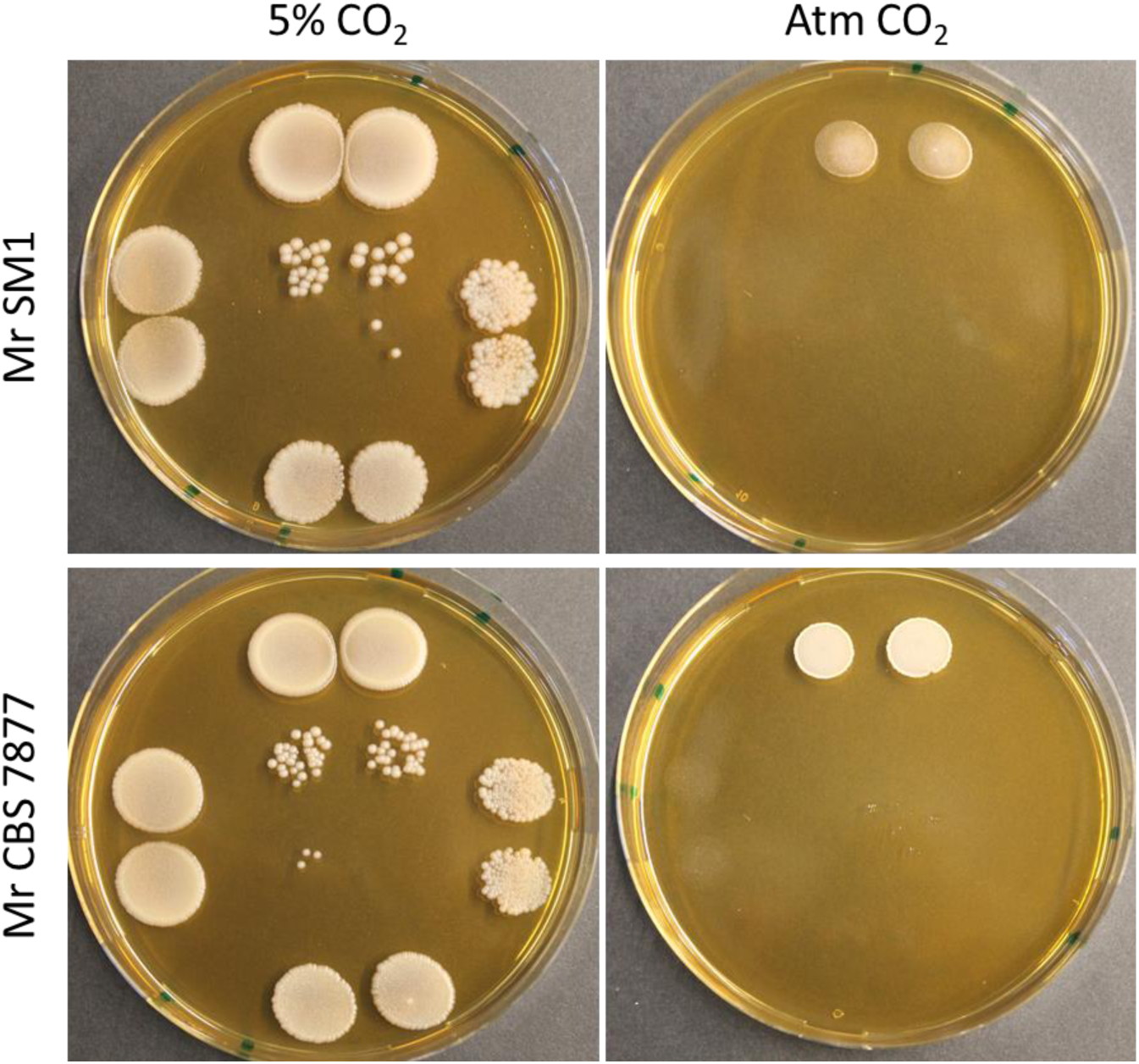
Supplementation of CO_2_ is necessary for *M. restricta* to form isolated colonies. Serial dilutions of *M. restricta* SM1 (scalp isolate) and *M. restricta* CBS 7877 (reference strain) cultures were spotted on mDixon plates and incubated under atmospheric levels of O_2_ with 5% CO_2_ supplementation or under atmospheric levels of both O_2_ and CO_2_. Undiluted spots are at the top and 10-fold dilutions were spotted counter-clockwise.

**Table 1.** Bulk growth characteristics of *M. restricta* (Mr)*, C. acnes* (Ca), and *S. epidermidis* (Se) strains under varying media and atmospheric conditions.

| Atmosphere <sup>a</sup> | Media | Mr (S) <sup>b</sup> | Mr (R) | Ca06 (S) | Ca15 (S) | Se (S) | Se (R) |
| --- | --- | --- | --- | --- | --- | --- | --- |
| Atm O<sub>2</sub><br>Atm CO<sub>2</sub> | mDixon | + <sup>c</sup> | + | - | - | + | + |
|  | 20 | - | - | - | - | + | + |
|  | TSA | - | - | - | - | + | + |
| 10% O<sub>2</sub><br>5% CO<sub>2</sub> | mDixon | + | + | - | - | + | + |
|  | 20 | - | - | - | - | + | + |
|  | TSA | - | - | - | - | + | + |
| 5% O<sub>2</sub><br>5% CO<sub>2</sub> | mDixon | + | + | - | - | + | + |
|  | 20 | - | - | - | - | + | + |
|  | TSA | - | - | - | - | + | + |
| 2% O<sub>2</sub><br>5% CO<sub>2</sub> | mDixon | + | + | - | - | + | + |
|  | 20 | - | - | + | + | + | + |
|  | TSA | - | - | + | + | + | + |
| Anaerobic<br>GasPak | mDixon | - | - | - | - | + | + |
|  | 20 | - | - | + | + | + | + |
|  | TSA | - | - | + | + | + | + |

<sup>a</sup> Plates were incubated at 32°C in a standard incubator with atmospheric levels of O<sub>2</sub> and CO<sub>2</sub>, a hypoxic chamber with indicated levels of O<sub>2</sub> and CO<sub>2</sub>, or an anaerobic container with GasPak with unknown levels of CO<sub>2</sub>.
<sup>b</sup> Scalp isolates are denoted as (S) and ATCC reference strains are denoted as (T).
<sup>c</sup> Cultures were streaked out on indicated plates and assessed after seven days as “+” for growth or “-” for no growth.

Based on this work, we next sought to identify a medium that supports the growth of all three species. The mDixon medium contains complex ingredients in addition to oleic acid and glycerol as defined carbon sources. mDixon supports the growth of *M. restricta* and *S. epidermidis*, but not *C. acnes* (Table 1), indicating that *C. acnes* requires additional nutrients. However, supplementing mDixon with simple sugars failed to grow *C. acnes* under our hypoxic condition (2% O_2_ and 5% CO_2_). We thus variably combined ingredients from Medium 20 and TSA to ultimately formulate a hybrid medium (D20), which reliably supports the growth of all three species as monoculture colonies (Fig. 2). We observe precipitates formed by *C. acnes* and *M. restricta* colonies, which are likely associated with the production of insoluble free fatty acids (29). In mixed-species colonies, however, *S. epidermidis* visibly dominated the population (Fig. S1), since *S. epidermidis* grows overnight while the other species take 5-7 days. We thus reduced the initial relative frequency of *S. epidermidis* by approximately a 1000-fold in mixed-species cultures to better enable the study of all three species in a single assay (Fig. 2) to use to study ecological interactions. Starting with a low frequency of *S. epidermidis* also better reflects the proportions observed on healthy scalp, where *S. epidermidis* is typically at lower frequency than *C. acnes* (10).

**Figure 2.**
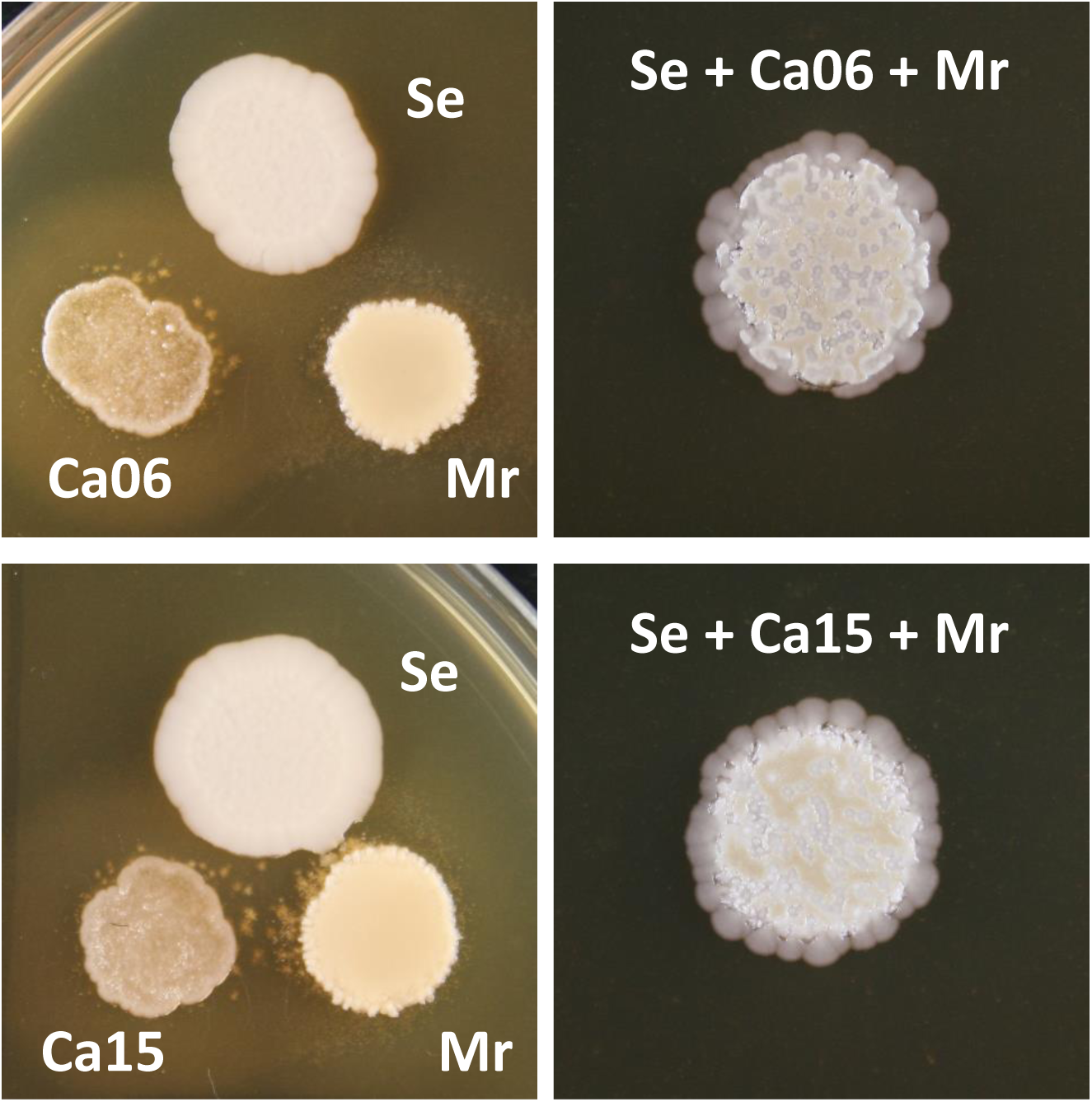
D20 is a new medium that commonly supports the growth of all three species. Scalp isolates of *S. epidermidis* (Se), *C. acnes* (Ca06 and Ca15), and *M. restricta* (Mr) grown as monoculture colonies (left) and mixed-species colonies (right). *S. epidermidis* was initially under-represented by approximately by a 1000-fold relative to *C. acnes* and *M. restricta* in the mixed-species colonies. The fuzzy zone surrounding *C. acnes* and *M. restricta* likely represent insoluble free fatty acids.

### Development of selective techniques for quantifying individual species in mixed populations

In order to experimentally study the ecology of scalp communities, one also needs a method to enumerating each species from co-cultures. *C. acnes* and *M. restricta* can be selected against by the presence or absence of O_2_, respectively, *C. acnes* does not grow on mDixon, and *M. restricta* fails to grow on Medium 20 or TSA. However, *S. epidermidis* grows on all media (Table 1). We therefore studied the effects of diverse antibiotics on *C. acnes* and *S. epidermidis* to identify candidates that exclusively select against *S. epidermidis*. *M. restricta* strains were excluded from this initial screen as they were not expected to be affected by the antibiotics. As summarized in Table 2, gentamicin, netilmicin, and tobramycin were uniquely more lethal to *S. epidermidis*.

**Table 2.** Antibiotic disc assay of *C. acnes* (Ca) and *S. epidermidis* (Se) strains.^a^.

| Antibiotic | Code | [ug] | Ca (R) | Ca06 (S) | Ca15 (S) | Se (R) | Se (S) |
| --- | --- | --- | --- | --- | --- | --- | --- |
| Amikacin | AK | 30 | 0.75 | 0.75 | 0.25 | 0.5 | 0.25 |
| Amoxycillin | AML | 25 | >3 | >3 | >3 | 1.25 | >3 |
| Amoxycillin-clavulanic acid | AMC | 30 | >3 | >3 | >3 | 1.75 | >3 |
| Aztreonam | APR | 30 | >3 | >3 | >3 | 0 | 0 |
| Cefepime | FEP | 30 | >3 | >3 | 2 | 2 | >3 |
| Ceftazidime | CAZ | 10 | 1.5 | 0.5 | 0.5 | 0.25 | 0.25 |
| Ciprofloxacin | CIP | 5 | 1.5 | >3 | 1.25 | 2 | 1.75 |
| Doripenem | DOR | 10 | >3 | >3 | 1.75 | >3 | >3 |
| Gentamicin | CN | 10 | 0.5 | 0.25 | 0.25 | 1 | 1.25 |
| Imipenem | IPM | 10 | >3 | >3 | >3 | >3 | >3 |
| Levofloxacin | LEV | 5 | >3 | 2.5 | 2 | 2 | 2 |
| Meropenem | MEM | 10 | >3 | 1.75 | 2 | >3 | >3 |
| Netilmicin | NET | 10 | 0.25 | 0 | 0 | 1 | 1.25 |
| Piperacillin | PRL | 30 | >3 | >3 | >3 | 1.25 | >3 |
| Piperacillin-tazobactam | TZP | 36 | >3 | >3 | >3 | 1.5 | >3 |
| Tobramycin | TOB | 10 | 0.25 | 0.25 | 0.25 | 0.75 | 0.75 |

<sup>a</sup> Discs impregnated with the indicated amount of the antibiotic [ug] were placed on a lawn of cells on TSA plates and the zone of no-growth was measured from the edge of each disc with respect to the diameter of each disc (e.g. 0.75 indicates approximately 75% of the length of the diameter from the edge of the disc to the growth boundary). >3 indicates zones that were more than 3 diameter distance away from the disc and 0 indicates the cultures grew at the edge of the disc. The longest recorded zone of no-growth for each antibiotic is shown above from assessing three independent discs after seven days of growth.

We next tested a broad concentration range of five readily available aminoglycosides (gentamicin, kanamycin, neomycin, streptomycin, and tobramycin) against all *C. acnes* and *S. epidermidis* strains. Since our ultimate goal was to identify an antibiotic concentration for exclusively counting isolated colonies of *C. acnes* from a mixture with *S. epidermidis*, we simply assessed the relative number of colonies from a serially diluted culture in the presence of the antibiotic to the no antibiotic control. Given the relatively large scale of this antibiotic screen, we initially assessed a single sample of each strain at each concentration and observed tobramycin at 8-32 ug/ml to be effective (Fig. S2). We next confirmed these results using triplicate samples against a narrower concentration range of tobramycin, and chose the concentration of 15 ug/mL for effectively selecting against *S. epidermidis* while not harming the scalp isolates of *C. acnes* (Fig. 3). In addition, *M. restricta* strains were unaffected at the same tobramycin concentration on mDixon as expected (Fig. 3).

**Figure 3.**
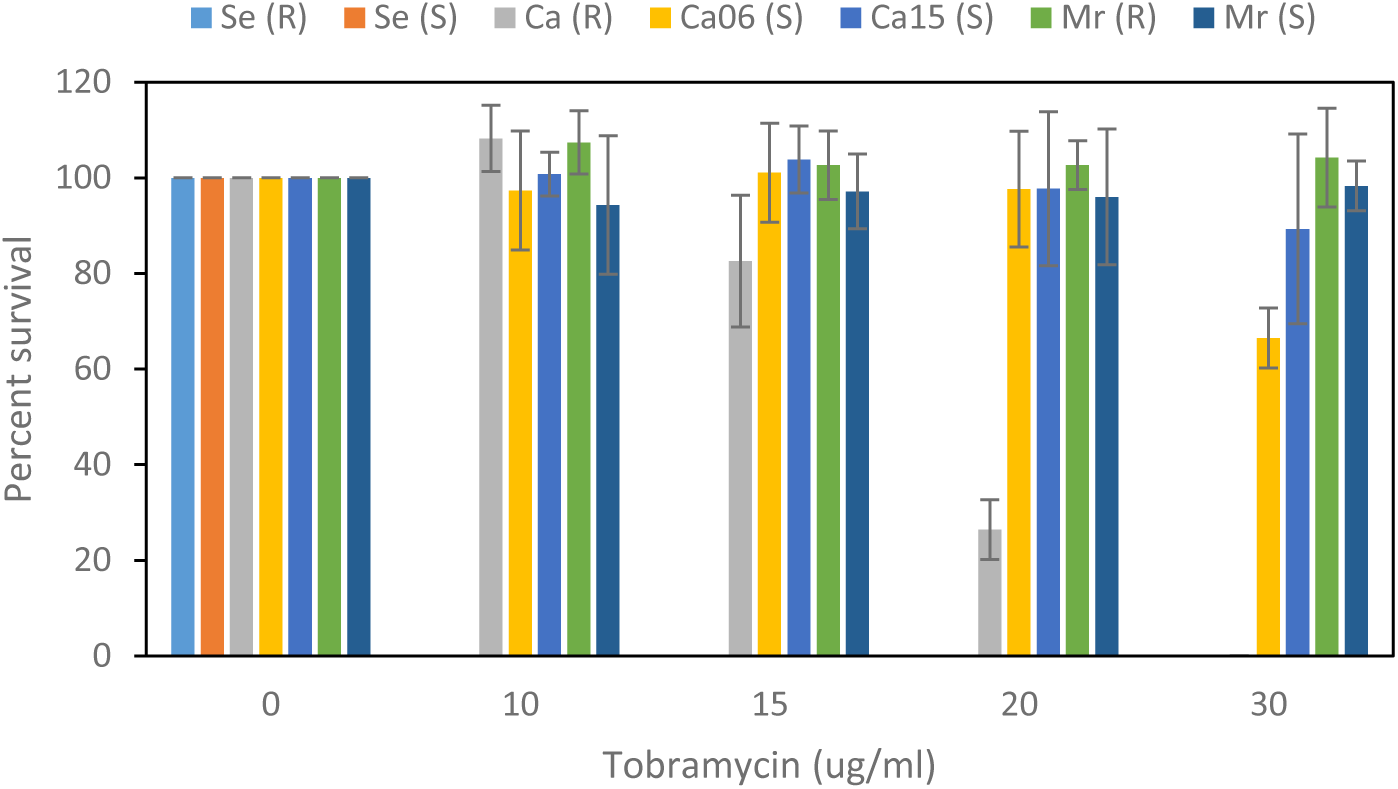
Tobramycin is selectively lethal against *S. epidermidis*. Serial dilutions of *S. epidermidis* (Se) and *C. acnes* (Ca) cultures were spotted on TSA plates containing different concentrations of tobramycin, and *M. restricta* (Mr) cultures were spotted on tobramycin-supplemented mDixon plates. (S) indicates scalp isolates and (R) indicates reference strains. Plotted are the mean of triplicate samples (antibiotic count over the no antibiotic control) and the error bars represent the standard deviation.

In summary, TSA supplemented with tobramycin at 15 ug/mL and incubated anaerobically exclusively selects for *C. acnes*, TSA incubated aerobically exclusively selects for *S. epidermidis*, and mDixon supplemented with tobramycin at 15 ug/ml and incubated aerobically with 5% CO_2_ exclusively selects for *M. restricta*. Furthermore, any spontaneously resistant colonies of *S. epidermidis* would be visible prior to the natural appearance of *C. acnes* or *M. restricta* colonies after 5-7 days of incubation.

### *C. acnes* exploits *M. restricta*, while *S. epidermidis* suppresses both *C. acnes* and *M. restricta* in mixed species culture mimicking healthy scalp

Having determined the common growth conditions for all three species (D20, 2% O_2_, 5% CO_2_, and 32°C) and the means to assess the abundance of each species in coculture, we next compared the growth dynamics of the scalp isolates in various assemblages over four time points. Day 7 was chosen as the earliest sampling time point since *C. acnes* and *M. restricta* require approximately seven days to exhibit appreciable colony growth in monoculture. Two scalp isolates of *C. acnes* strains were included in the assays for robustness, and also because a reduction in the relative load of *C. acnes* appears to be a conserved signature for poor scalp health (11–13). We calculated the relative growth (*r*) of each focal strain in two-species or three-species cultures compared to monoculture, which represents natural log increase (positive values indicate advantage) or decrease (negative values indicate disadvantage) in population size. The two *C. acnes* strains produced similar patterns, where they both benefit from the presence of *M. restricta*, but are negatively affected by *S. epidermidis* (Fig. 4A-B). *C. acnes* is negatively affected by the presence of both *M. restricta* and *S. epidermidis*, but this is largely due to the impact of *S. epidermidis* alone. In contrast, *S. epidermidis* appears to be mildly suppressed by *M. restricta* at earlier time points, but it is largely unaffected by the presence of either or both *C. acnes* and *M. restricta* (Fig. 4C). *M. restricta* is suppressed by the presence of *S. epidermidis* in both two-species and three-species mixtures, and it appears to be negatively impacted by *C. acnes* as indicated by the consistently negative relative growth values (Fig. 4D). Overall, therefore, we find evidence of strong and diverse pairwise interactions among the scalp microbes, which appear to be overshadowed by the presence of *S. epidermidis* in three-species mixtures. Furthermore, fitting linear models to the growth dynamics of each strain revealed the same patterns of pairwise interactions (Fig. S3).

**Figure 4.**
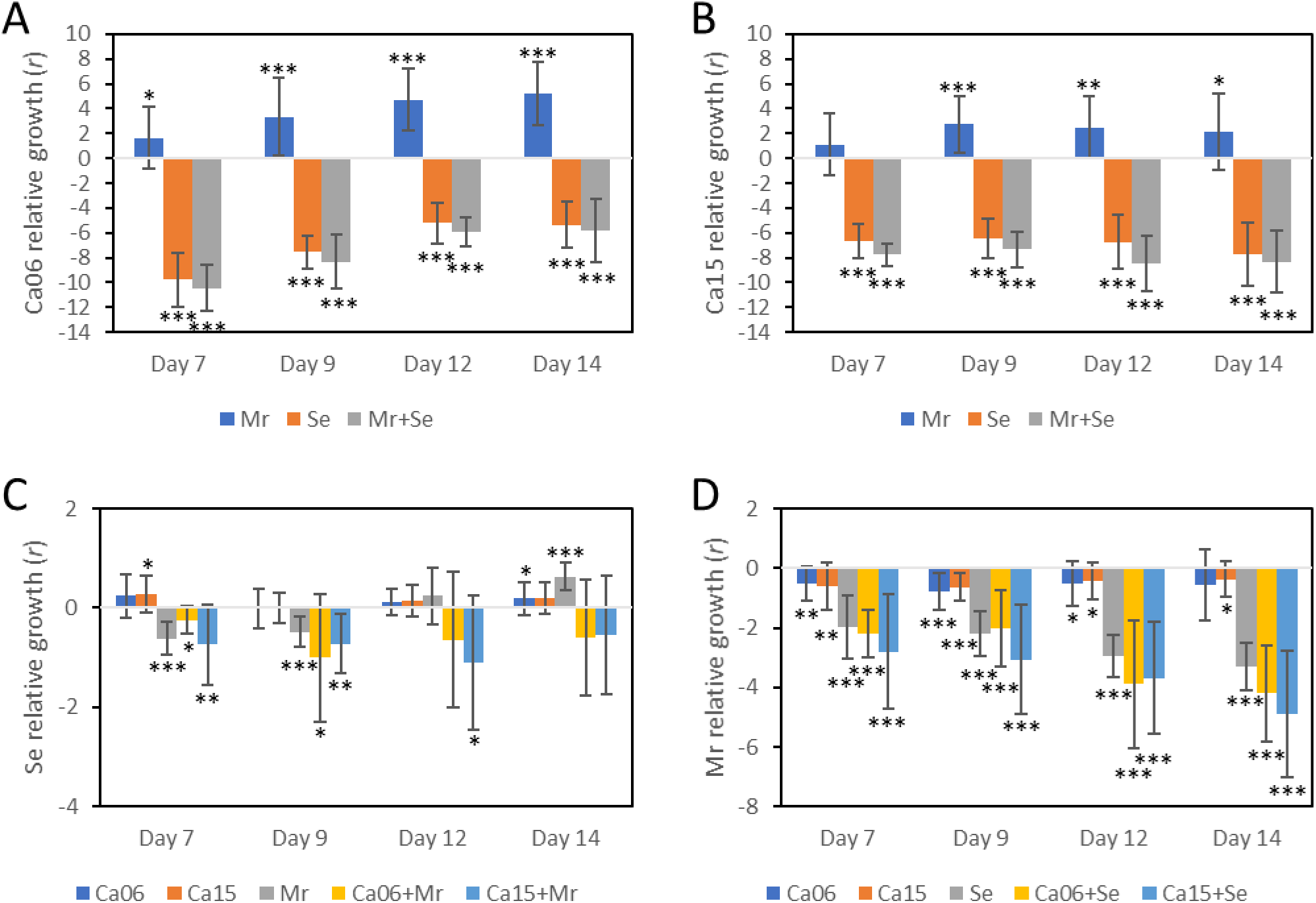
Co-culture growth captures beneficial, antagonistic, and neutral interactions among the scalp isolates. Shown are the mean relative growth of *C. acnes* 06 (A), *C. acnes* 15 (B), *S. epidermidis* (C), or *M. restricta* (D) in two-species or three-species cultures compared to the respective monoculture. Specific strains that were mixed with the focal strain are indicated at the bottom of each panel. The y-axis represents natural log increase (positive values indicate advantage) or decrease (negative values indicate disadvantage) in population size. A single-sample student’s t-test was conducted to determine the significance of difference between the observed mean and the hypothetical mean of zero. * represents two-tailed p < 0.05, ** represents p < 0.01, and *** represents p < 0.001; n = 16 (*C. acnes* and *M. restricta* pairwise co-cultures) and n=12 (all other co-cultures); error bars represent the standard deviation of the mean.

## Conclusion

We have overcome significant technical challenges to develop the first experimental model system to empirically assess interactions among the dominant bacteria and fungus of the human scalp. Our assay does not include all species that exist on the scalp. For example, *M. globosa* can be a significant fungal member of the scalp microbiota in certain human populations (30–32). Nevertheless, our work allows us to experimentally study the ecological interactions among the three dominant members of the scalp for the first time. This work reveals the potential for strong ecological interactions (Fig. 5). Notably *C. acnes* benefits from *M. restricta,* but harms it in return, and that *S. epidermidis* harms both other species. Moreover, when all three species are cultured, it is the effects of *S. epidermidis* that dominates and suppresses the growth of both others.

**Figure 5.**
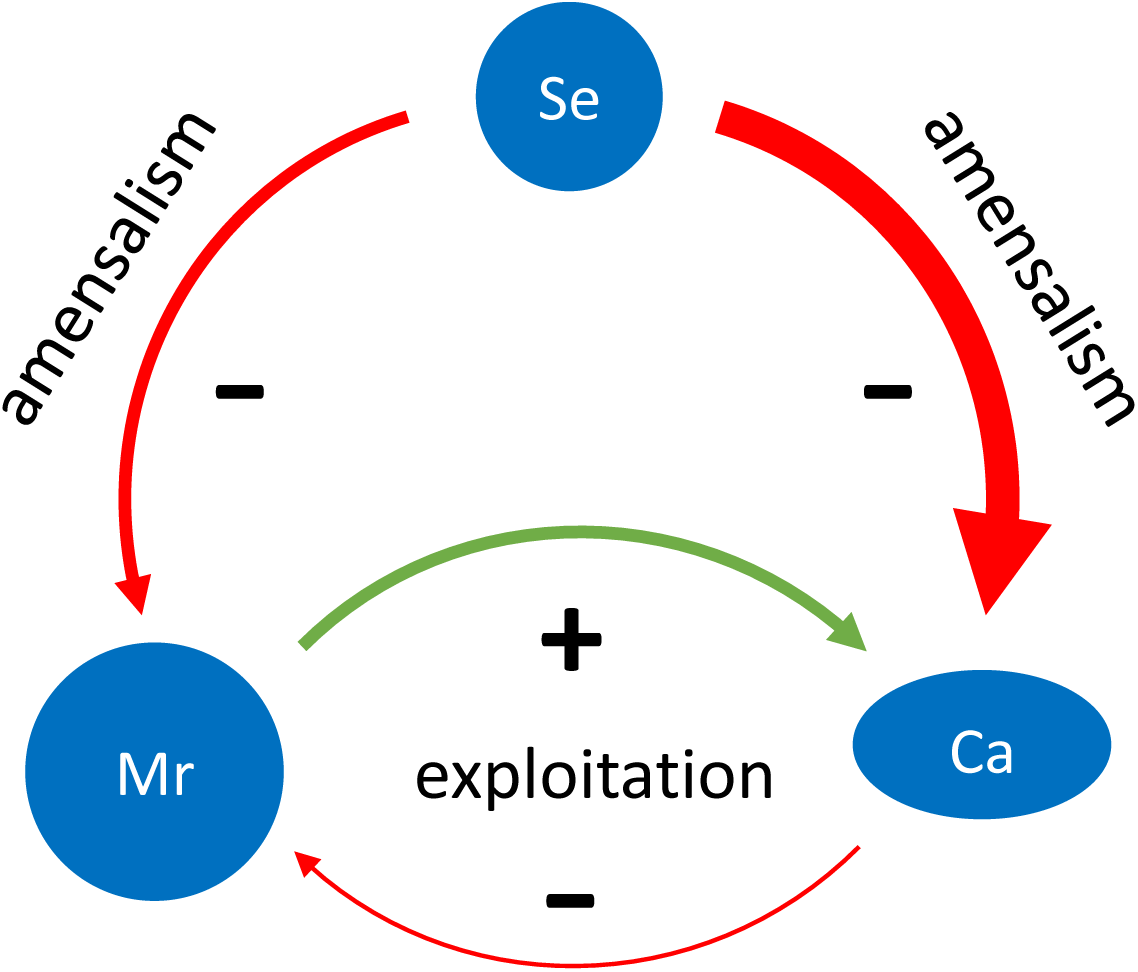
Ecological network of scalp microbes. *S. epidermidis* suppresses both *C. acnes* and *M. restricta*, but the impact is dramatically stronger against C. acnes. In contrast, S. epidermidis neither benefits from nor is harmed by the presence of M. restrica and/or C. acnes (amensalism). C. acnes greatly benefits from the presence of M. restricta, but moderately harms it in return (exploitation).

An important caveat to this last observation is that, in our assay, *S. epidermidis* has the ability to overgrow both *C. acnes* and *M. restricta*, which is not seen in metagenomics data, even though *S. epidermidis* abundance is higher in dandruff versus healthy scalp (11–13). There are differences between natural conditions and our assay that suppress the domination of *S. epidermidis*: such as the absence of corneocytes that supports the adhesion of *S. epidermidis* (33) and the absence of skin antimicrobial peptides (34). Nevertheless, it is interesting to compare our findings to the data on community composition during seborrehic dermatitis and dandruff. Here, sequencing work has found that *S. epidermidis* and *M. restricta* increase, while *C. acnes* declines (12, 13). These patterns suggest that *M. restricta* is somehow able to avoid the negative effects of *S. epidermidis* that we observe here. This ability could lie in differences in chemical conditions *in vivo* that weaken this particular interaction, or spatiogenetic structure that keeps the species apart. While the negative interaction between *S. epidermidis* and *M. restricta* is not seen in the metagenomics data, the inverse shift seen in the two bacteria during seborrehic dermatitis and dandruff is consistent with the strong negative effect of *S. epidermidis* that we observe on *C. acnes*. Indeed, the shift in the bacteria is the most robust pattern found with the onset of dandruff; some studies find no change in *M. restricta* (11). Altogether, these observations suggest a potential role for bacterial community composition in dandruff, and raise the possibility of treatment strategies that seek to promote the growth of *C. acnes* over *S. epidermidis* (11–13, 35).

## Methods

### Strains and culture conditions

*S. epidermidis* V17610, *C. acnes* OA006 (IA-2), *C. acnes* OA0015 (IV), and *M. restricta* SM1 were isolated from the human scalp and provided by L’Oréal Research and Innovation. *S. epidermidis* ATCC 12228, *C. acnes* ATCC 6919 (IA-2), and *M. restricta* CBS 7877 are reference type strains provided by L’Oréal Research and Innovation. Tryptic soy broth (TSB; Fisher) was used for liquid cultures of *S. epidermidis* and *C. acnes* strains, and modified Dixon (mDixon) (36) was used for liquid cultures of *M. restricta* strains. Medium 20 (37) was also used as specified. TSB, mDixon, and Medium 20 were solidified with agar (15 g/L) to prepare plates, and solidified TSB is denoted as TSA. D20 [15 g tryptone (Difco), 10 g yeast extract (Difco), 0.50 g cysteine hydrochloride (Fisher), 6 g mycological peptone (Oxoid), 18 g malt extract (Oxoid), 5 g oxgall (Difco), 12 g agar (Fischer), 2 mL glycerol (Fisher), 2 mL oleic acid (Fisher), 10 mL tween-40 (Sigmal Aldrich), made up to 1 L with deionized water] was autoclaved then supplemented with 50 mL 10% mannitol (Fisher) and 25 mL hemin solution [0.1 g hemin chloride (Roth), 4 mL triethanolamine (Prolabo), 96 mL deionized water, filter-sterilized]. When required, antibiotics (Sigma Aldrich) were added at indicated concentrations. *M. restricta* liquid cultures were incubated at 30°C with shaking at 250 rpm and plate cultures were sealed with gas-permeable parafilm and incubated at 32°C with or without 5% CO_2_ as indicated. *S. epidermidis* was grown at 37°C with shaking at 250 rpm for liquid cultures. *C. acnes* was grown anaerobically at 37°C in the BD GasPak EZ system with the BD anaerobic indicator strip, and liquid cultures were manually shaken once a day. All lab work was conducted in a biosafety laminar flow hood to minimize contaminations.

The following protocols were used to prepare liquid cultures for all experiments. *M. restricta* frozen stocks were thawed at room temperature and 20 ul was spotted on mDixon plates and incubated for five days. Cells were then scraped with a sterile loop to inoculate 20 mL mDixon and incubated for five days. *C. acnes* frozen stocks were streaked out on TSA plates, incubated for seven days, 30 isolated colonies inoculated into 10 mL TSB with a porous cover to permit gas exchange, and incubated for seven days with manual shaking once a day to displace any sunken aggregates. *S. epidermidis* frozen stocks were streaked out on TSA plates, incubated overnight, a single isolated colony inoculated into 5 mL TSB, and incubated overnight.

### Antibiotic disc assay

Liquid cultures (150 uL) of *S. epidermidis* and *C. acnes* were spread out on TSA plates with a sterile bent glass rod and three antibiotic discs (Oxoid) for each antibiotic were placed in a triangular format with equal distance. Plates were incubated for seven days and the zone of no growth from the edge of the disc was estimated with respect to the diameter of each disc.

### Aminoglycoside assay

Liquid cultures of *S. epidermidis* and *C. acnes* were serially diluted in sterile phosphate buffered saline (PBS) and two 50 ul aliquots of each dilution was spotted on TSA plates and TSA plates with gentamicin, kanamycin, neomycin, streptomycin, or tobramycin at the indicated concentrations. Plates were incubated for seven days and isolated colonies were counted. Percent survival was calculated by comparing the CFU on antibiotic plates to CFU on the control plates without the antibiotic. Same methods were used for the narrower concentration range assay with tobramycin, except that the samples were assessed in triplicate.

### Monoculture and co-culture experiments

Liquid cultures were sonicated in 5 mL volumes in a 50 mL tube at 20% amplitude (500W) for 20 seconds (with cycles of 1 second on and 1 second off) to displace any aggregates of cells. We established that sonication at this low energy setting consistently yielded higher CFU counts for all strains compared to no sonication, indicating that the sonication condition does not harm the cells, and we further confirmed the effective generation of individual cells by microscopy. *S. epidermidis* was serially diluted in PBS to 10^-5^, which represents approximately 1000-fold reduction in density compared to undiluted *C. acnes* and *M. restricta* samples. Using the diluted liquid culture of *S. epidermidis* and undiluted liquid cultures of *C. acnes* or *M. restricta*, three-species mixtures were prepared by mixing 200 uL of each strain, two-species mixtures were prepared by mixing 200 uL of each strain with 200 uL PBS, and monoculture mixtures were prepared by mixing 200 uL of each strain with 400 uL PBS. 10 uL of each mixture was spotted three times on D20 plates in a triangular format with equal distance. Once the spots dried, plates were wrapped in gas-permeable parafilm and incubated in the Baker-Ruskinn Invivo2 Plus chamber set at 2% O_2_, 5% CO_2_, and 32°C. Each mixture was also serially diluted in PBS and two 20 uL aliquots of each dilution was spotted on TSA (*S. epidermidis*), TSA with 15 ug/mL tobramycin (*C. acnes*), and mDixon with 15 ug/mL tobramycin (*M. restricta*). *S. epidermidis* plates were counted after one day and *C. acnes* and *M. restricta* plates were counted after seven days to calculate the CFU of each species initially spotted on the D20 plates (day 0). On days 7, 9, 12, and 14, monoculture and co-culture colonies were scraped using a sterile bent Pasteur pipette and the cells were suspended in 5 mL PBS, vortexed, sonicated, serially diluted in PBS, and spotted on selection plates, incubated, and CFU calculated as described above. All experiments were conducted with a minimum of 12 biological replicates. The relative growth of each focal strain in two-species and three-species mixtures were compared to monoculture growth by calculating the selection rate (*r*) (38).

### Statistical analysis

A single-sample student’s t-test was conducted in GraphPad Prism to determine the significance of difference between the observed mean relative growth (*r*) and the hypothetical mean of zero. To examine the potential effect of each focal strain’s growth by the presence of other species, individual linear regressions were fit on the abundances of each strain from day seven onwards. In each case, the response variable was the logged CFU of the focal strain, while the dependent variables were the logged CFUs of each of the other strains and day. Models were fit using the lme4 package in R (version 4.0.2).

### Data availability

The raw data from monoculture and co-culture experiments are available at https://github.com/wook-kim-lab/scalp-microbiota-model.

## Acknowledgements

We thank B. Balasubramonian, J. Chen, S. Dighole, M. Echard, T. Cupac, D. Jungo, and S. Li for technical assistance. This study was funded by L’Oréal Research and Innovation C140918 (K.R.F. and W.K.) and the Hunkele Dreaded Disease Award G1800049 (W.K.). K.Z.C. is supported by a University of Manchester Presidential Fellowship. K.R.F. is supported by European Research Council Grant 787932, and by Wellcome Trust Investigator award 209397/Z/17/Z. K.R.F. and W.K. designed the study and analyzed data; W.K. wrote the manuscript; K.Z.C., C.C., K.R.F., and W.K. edited the manuscript; S.B., T.R., S.C., A.S., W.M., and W.K. performed experiments; K.Z.C. performed mathematical modeling and analysis; C.C. provided strains, reagents, and advice.

## Supplemental Material

Figs S1 to S3

---

**Figure S1.**
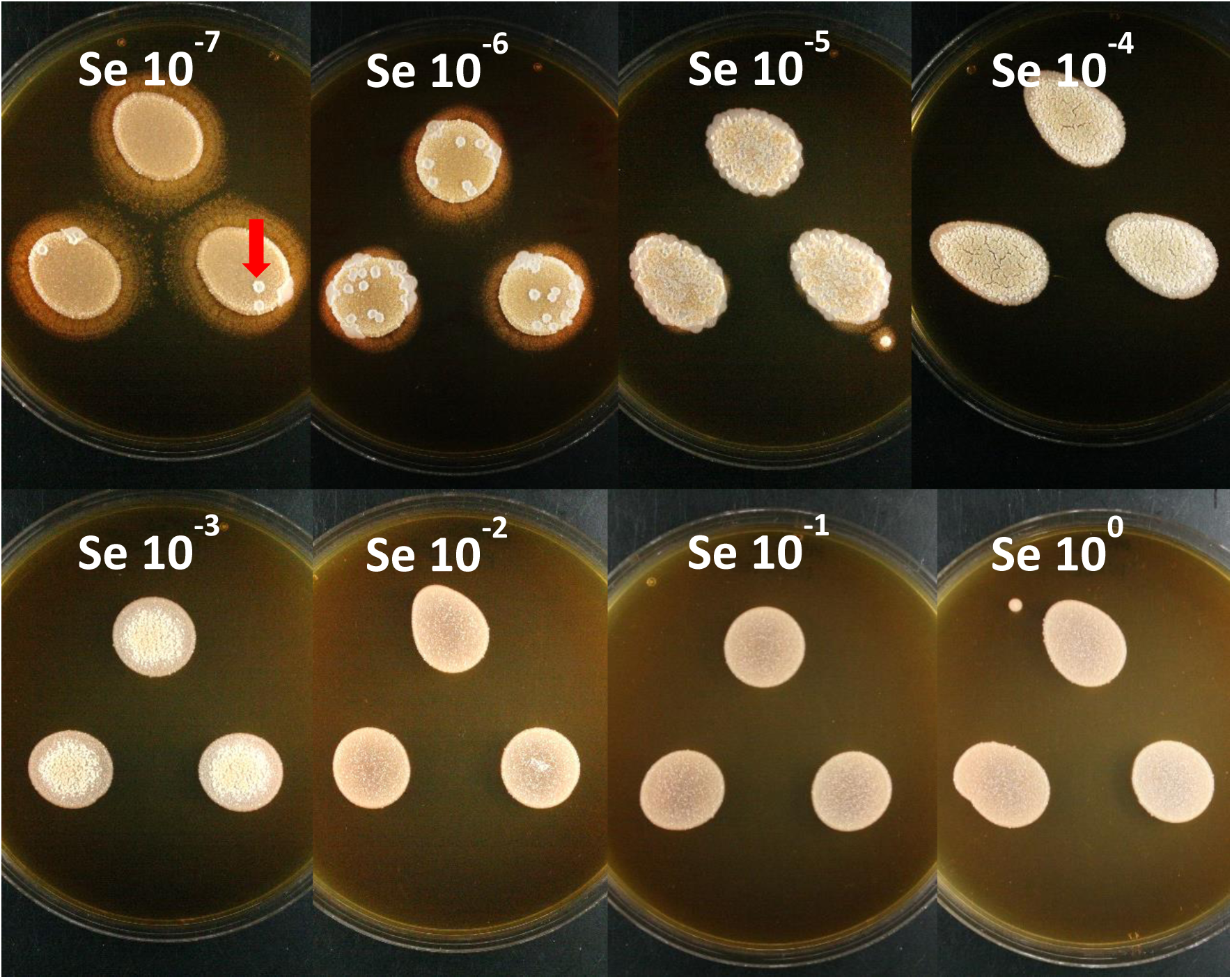
*S. epidermidis* overtakes co-cultures unless it is initially under-represented. Each panel shows three replicate co-culture colonies resulting from a mixture that initially contained equal proportions of *M. restricta* and *C. acnes* with serial dilutions of *S. epidermidis* (reference strain). 10^0^ indicates undiluted *S. epidermidis*. Note the individual colonies of *S. epidermidis* at 10^-6^ and 10^-7^ (red arrow).

**Figure S2.**
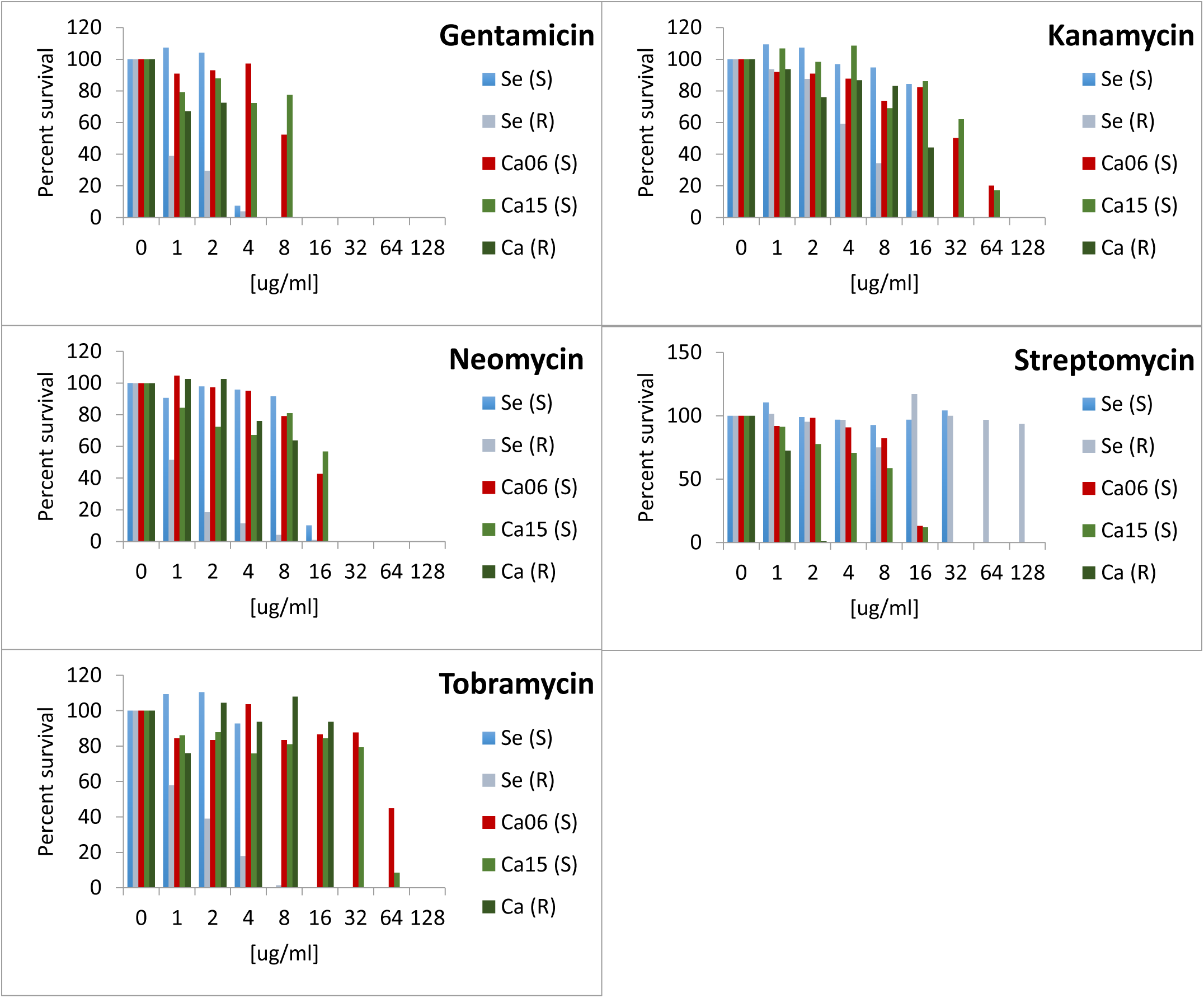
*S. epidermidis* and *C.* acnes strains exhibit varying levels of sensitivity to aminoglycoside antibiotics. Serial dilutions of *S. epidermidis* (Se) and *C. acnes* (Ca) cultures were spotted on TSA plates containing different concentrations of the indicated antibiotic across a broad range (S) indicates scalp isolates and (R) indicates reference strains. Plotted is the antibiotic count over the no antibiotic control of a single sample.

**Figure S3.**
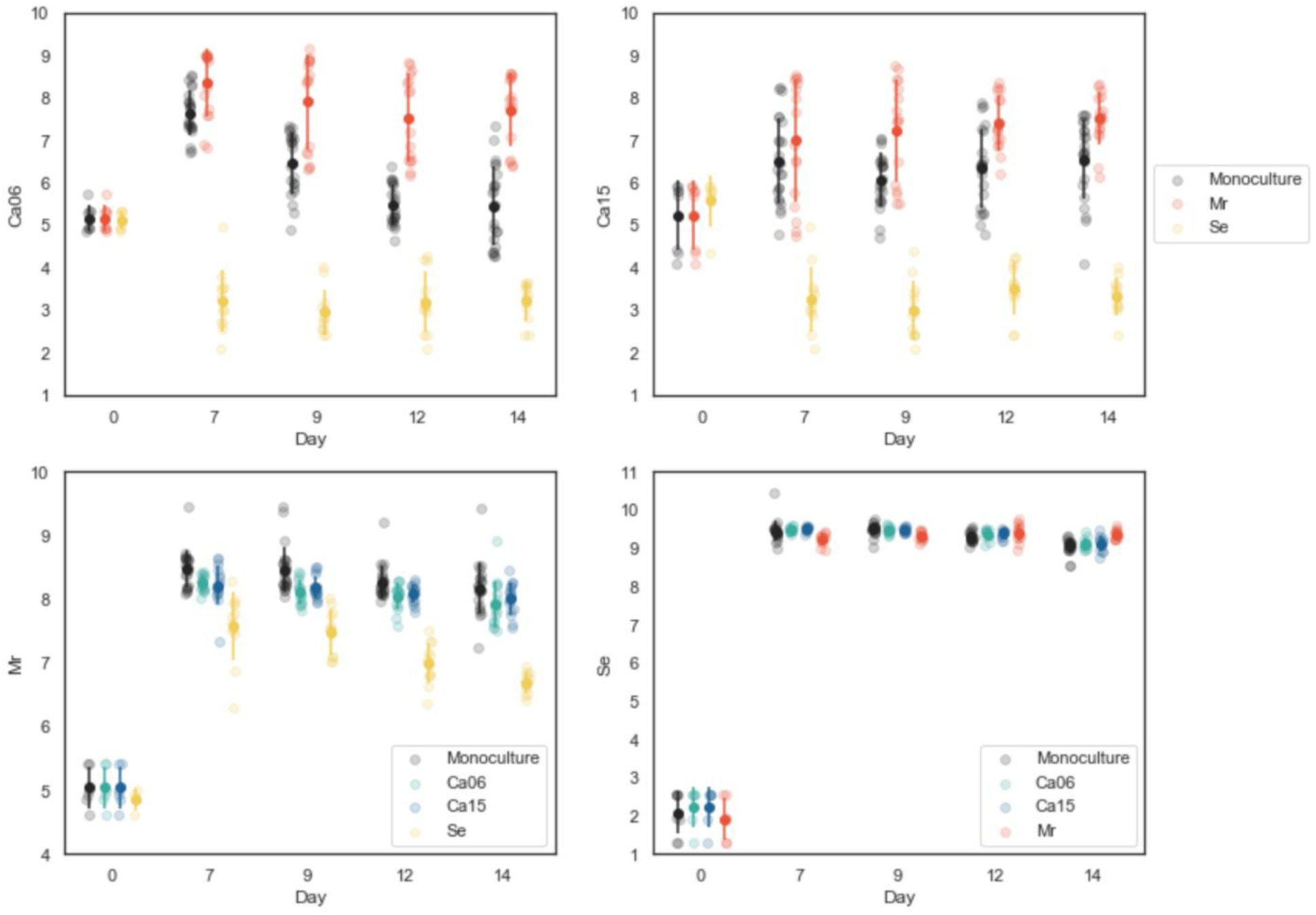
Comparisons of raw CFU counts of monoculture and paired co-cultures capture diverse interactions. The focal strain in each plot is indicated on the left and the co-presence of specific species is indicated in the figure legend. The y-axis represents the log_10_CFU values and plotted are independent samples. Fitting linear models to the growth dynamics of each strain revealed ecological interactions between the species. Both *C. acnes* (Ca) strains 06 and 15 exhibited enhanced growth in the presence of *M. restricta* (Mr) (effect size 0.20 and 0.11 respectively, p <0.001 in each case), but reduced growth in the presence of *S. epidermidis* (Se) (effect size -0.33 and p<0.001 for both). *S. epidermidis* did not show any significant difference in growth when cultured with any of the other strains (p >0.1 for each). *M. restricta* showed slightly reduced growth in the presence of the *C. acnes* strains (effect size -0.03 and p <0.001 for both) and reduced growth in the presence of *S. epidermidis* (effect size -0.12, p <0.001).

